# Synthesis and characterization of peptide conjugated human serum albumin nanoparticles for targeted cardiac uptake and drug delivery

**DOI:** 10.1101/2021.07.23.453538

**Authors:** Nikita Lomis, Susan Westfall, Dominique Shum-Tim, Satya Prakash

**Affiliations:** Biomedical Technology and Cell Therapy Research Laboratory, Department of Biomedical Engineering, 3775 University Street, Montreal, QC, H3A 2B4, Canada; Division of Experimental Medicine, 1001 Boulevard Décarie, Montréal, QC, H4A 3J1, Canada; Department of Neurology, Icahn School of Medicine Mount Sinai, New York, NY, 10029, USA; Division of Cardiac Surgery and Surgical Research, Royal Victoria Hospital, 1001 Boulevard Décarie, Montréal, QC, H4A 3J1, Canada

**Keywords:** Albumin, drug delivery, nanoparticles, peptide, angiotensin, heart failure

## Abstract

Congestive heart failure, a prominent cardiovascular disease results primarily from myocardial infarction or ischemia. Milrinone (MRN), a widely used clinical drug for heart failure, improves myocardial contractility and cardiac function through its inotropic and vasodilatory effects. However, lacking target specificty, it exhibits low bioavailability and lower body retention time. Therefore, in this study, angiotensin II (AT1) peptide conjugated human serum albumin nanoparticles (AT1-HSA-MRN-NPs) have been synthesized for targeted delivery of MRN to the myocardium, overexpressing AT1 receptors under heart failure. The NPs were surface functionalized through a covalent conjugation reaction between HSA and AT1. Nanoparticle size was 215.2±4.7 nm and zeta potential -28.8±2.7 mV and cumulative release of MRN was ∼72% over 24 hrs. The intracellular uptake of nanoparticles and cell viability was studied in H9c2 cells treated with AT1-MRN-HSA-NPs vs the control non-targeted drug, MRN Lactate under normal, hypoxic and hypertrophic conditions. The uptake of AT1-HSA-MRN-NPs in H9c2 cells was significantly higher as compared to non-targeted nanoparticles, and the viability of H9c2 cells treated with AT1-MRN-HSA-NPs vs MRN Lactate was 73.4±1.4% vs 44.9±1.4%, respectively. Therefore, AT1-HSA-MRN-NPs are safe for *in vivo* use and exhibit superior targeting and drug delivery characteristics for treatment of heart failure.

## Introduction

Cardiovascular disease (CVD) is one of the leading causes of mortality, with incidences constantly on the rise across the developed and developing world. Of these CVDs, myocardial infarction (MI) and congestive heart failure (CHF) are responsible for more than 50% of the global cases of CVDs with high rates of readmission and re-hospitalization ^1^. Currently, the most common treatments for CHF include surgical interventions such as heart transplant, stenting, bypass surgeries, ventricular assist devices and medical treatments including drugs such as ACE inhibitors, beta blockers, vasodilators etc. ^2,3^. However, despite these common treatment modalities, the average survival rate for more than 50% of patients who have suffered a first heart failure is less than 5 years ^4^. Therefore, there is an urgent need for development of effective therapies to target these diseases.

CHF is typically caused by blockage of the coronary artery, which results in ischemia and leads to an irreversible necrosis of the cardiomyocytes. It is widely suggested that under MI and CHF, an overexpression of the angiotensin II type 1 receptors (AT1Rs) on the myocardium may be an underlying cause for cardiac remodeling ^5-8^. This facilitates the underlying mechanism of ACE inhibitor drugs, which block the overexpressed AT1 receptors to prevent cardiac hypertrophy ^9,10^. The condition of overexpression of AT1Rs has been explored by a study in which superior AT1R targeting was reportedly achieved for delivery of AT1 bound quantum dot nanoparticles ^11^. The use of AT1-conjugated liposomes in a mouse model of MI has also shown their specific internalization ^5^. This will be the first study demonstrating a new synthesis scheme for development of a novel nanoparticle formulation, AT1-HSA-MRN-NPs, for targeted drug delivery to the heart.

Previously, we presented the preparation and binding of HSA-NPs with milrinone (MRN), a cardiac inotrope and vasodilator drug, widely used for the treatment of CHF ^12^. Milrinone is a phosphodiesterase III enzyme inhibitor which increases the intracellular cAMP concentration, providing higher calcium influx to create a positive inotropic effect ^13-15^. Clinically administered as a lactate formulation (MRN Lactate), milrinone is known to improve the overall cardiac function by increasing myocardial contractility and decreases systemic vascular resistance ^16,17^. However, the lack of target specificity, lower bioavailability and a half-life of approximately 1-2 hours in humans, necessitates its use as a continuous infusion, also causing other side effects such as renal dysfunction, palpitations and arrhythmias ^18,19^.

The enhanced targeting and delivery of MRN could be achieved by loading it on nanoparticles targeted to the intended site of action. One of the most widely used nanoparticles are HSA-NPs owing to unique features like biocompatibility, biodegradability and non-immunogenicity. The HSA molecule possesses multiple pockets to promote binding of various hydrophilic and hydrophobic drugs such as paclitaxel, doxorubicin etc. ^20-22^. Computational modeling and enzyme release studies have shown that MRN binds hydrophobically to the HSA molecule ^23^. HSA-NPs are known to improve the blood circulation time of otherwise insoluble or free drugs and are anticipated to improve the bioavailability of MRN as well ^24,25^.

The presence of active functional groups on HSA allows opportunities for surface-modification to bind additional ligands such as peptides, antibodies, genes, and other molecules ^26-29^. This would be useful in enhancing receptor mediated nanoparticle internalization and drug delivery characteristics. In this study, HSA was surface modified through a two-step reaction scheme to bind the AT1 peptide to form AT1-HSA. The AT1-HSA was further bound to the cardiac inotrope and vasodilator drug, MRN, to form AT1-HSA-MRN-NPs. In addition to studying drug binding and release, cell viability analysis was performed to compare the safety of AT1-HSA-MRN-NPs with MRN Lactate, which is the clinical drug formulation for CHF treatment. These studies indicate that the novel AT1-HSA-MRN-NPs can bind and deliver MRN and must be tested *in vivo* for use in cardiovascular diseases.

## Materials and Methods

### Materials

Human serum albumin (> 97% lyophilized) was purchased from Sigma Aldrich (Oakville, ON, Canada). Glutaraldehyde (25% aq. solution) was purchased from Alfa Aesar (Cedarlane, Burlington, ON, Canada). Fluorescein isothiocynate human serum albumin (FITC-HSA) was purchased from Sigma Aldrich (Oakville, ON, Canada). Milrinone was purchased from Selleck Chemicals (Burlington, ON, Canada). Bradford reagent was purchased from Bio-Rad (St. Laurent, QC, Canada). The 5(6)-Carboxyfluorescein *N*-hydroxysuccinimide was purchased from Thermo Fisher Scientific (ON, Canada) Other chemicals were purchased from Fisher Scientific (Nepean, ON, Canada).

### Synthesis of AT1-peptide

The Angiotensin II Type 1 (AT1) receptor targeting peptide is a chain of 8 amino acids Asp-Arg-Val-Tyr-Ile-His-Pro-Phe, and was synthesized by CanPeptide (Pointe-Claire, QC, Canada) as NH_2_-Gly-Gly-Gly-Gly-Asp-Arg-Val-Tyr-Ile-His-Pro-Phe-NH_2_ along with a Scrambled Peptide NH_2_-Gly-Gly-Gly-Gly-Phe-His-Tyr-Arg-Asp-Val-Ile-Pro-NH_2_ following the sequence mentioned by Dvir et al.^5^.

### Surface modification of HSA with AT1 peptide

The surface of HSA was modified to facilitate attachment of the AT1 peptide in a two-step reaction. An aqueous solution of 20 mg/mL of HSA dissolved in deionized water (0.3 mM) was prepared and reacted with a 10-fold molar excess of 5(6)-Carboxyfluorescein *N*-hydroxysuccinimide ester (COOH-FITC-NHS)), for 1 hour. This was further reacted with EDC/Sufo-NHS for 30 minutes followed by reaction with either the AT1 or scrambled peptide (Scr) for 4 hours **(Fig. 1)**. The AT1 and Scr peptide was added in a 10-fold molar excess than HSA. The AT1-HSA and Scr-HSA were purified by dialysis using the Slide-a-Lyze dialysis cassette (10,000 Da MWCO). The purified sample was then lyophilized and stored at 4°C.

**Figure 1.**
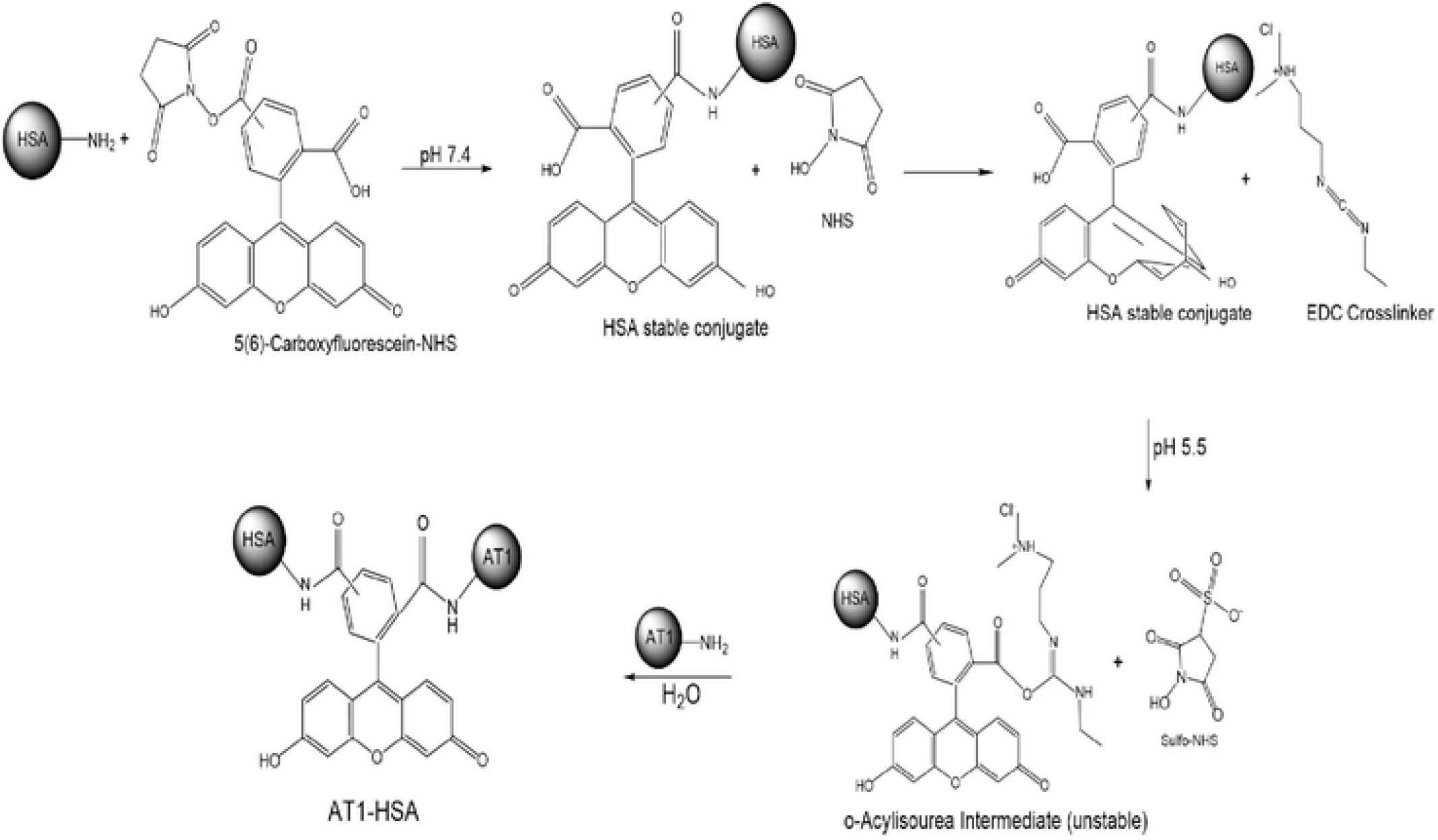
Schematic representation of the HSA surface modification to bind the AT1 peptide through a two-step chemical conjugation reaction using heterobifunctional cross-linkers.

### Nuclear Magnetic Resonance

1D proton spectra were recorded at 25° on a 500 MHz Varian INOVA NMR Spectrometer, with an HCN triple resonance RT probe with Z-axis pulsed field gradients. Spectra were recorded with double pulsed-field gradient spin echo for suppression of residual H_2_O signal. 256 scans with a recycle delay of 1s were collected with a sweep width of 8000 Hz and an acquisition time of 1s. Data were processed with 1 Hz line broadening using the VNMRJ 4.2 software. The concentrations of HSA, AT1 peptide and AT1-HSA were 5 mg/mL in D_2_O.

### Mass Spectrometry

The AT1-HSA, HSA and AT1 peptide samples were analyzed by the Matrix-Assisted Laser Desorption/Ionization Time of Flight Mass Spectrometry (MALDI-TOF-MS) MALDI Autoflex III-TOF-(BRUKER) SMARTBEAM) (Dept. of Chemistry, McGill University, Montreal, QC, Canada) in linear positive mode. Dihydroxybenzoic Acid was used as the Matrix and the AT1-HSA, HSA and AT1 peptide samples were dissolved in water at concentrations of approximately 7 mg/mL.

### Nanoparticle Preparation

The AT1-HSA-MRN-NPs were prepared by following the ethanol desolvation technique ^30,31^. Briefly, an aqueous solution of 20 mg/mL of AT1-HSA was prepared in deionized water. The pH of the solution was adjusted to pH 8.0 using 0.1 M NaOH. MRN was dissolved in DMSO and added to the AT1-HSA solution with a final MRN/HSA ratio of 1:20 by weight. Ethanol was added in a dropwise manner resulting in solution turbidity. Glutaraldehyde (8% v/v aq. solution) was added to the reaction mixture at a concentration of 0.588 μl/mg HSA and polymerized for 24 hours. AT1-HSA was substituted with Scr-HSA for preparation of Scr-HSA-NPs and with FITC-HSA for preparation of FITC-HSA-MRN-NPs. The nanoparticles were washed by three rounds of ultracentrifugation at 16500 rpm for 15 minutes each at 25°C. Supernatant was collected for detection of unbound MRN. The pellet was washed with deionized water and finally re-dispersed in phosphate buffer saline (PBS). The nanoparticles were tip-sonicated for 10 minutes and stored at 4°C.

### Nanoparticle characterization, yield and encapsulation efficiency

The average particle size of the nanoparticles was measured by Dynamic Light Scattering (DLS) using a Particle Size Analyzer (Brookhavens Instruments Corporation, NY, USA). The samples were diluted 1:20 with deionized water and measured at a scattering angle of 90° and temperature of 25 °C. The Polydispersity Index (PDI) estimated the size distribution of the nanoparticles. The zeta potential was measured by a Zeta Potential Analyzer (Brookhavens Instruments Corporation, NY, USA) using electrophoretic laser Doppler anemometry. The size and shape of the nanoparticles were examined by Transmission Electron Microscopy (TEM) (FEI Tecnai G^2^ Spirit Twin 120 kV Cryo-TEM, Gatan Ultrascan 4000 4k x 4k CCD Camera System Model 895).

The yield of the nanoparticles was measured by the UV-spectrophotometric method. A standard curve of HSA solution dissolved in Bradford reagent was used as a reference and absorbance was measured at 595 nm. To determine encapsulation efficiency, nanoparticles were spin concentrated using centrifugal filters with molecular weight cut off (MWCO) of 10,000 Da for eluting the non-encapsulated MRN into the collection tube. A standard curve of MRN in a mixture containing DDQ/Ethanol was used as a reference and absorbance was measured at 356 nm ^32^.

### *In Vitro* Milrinone Release from Nanoparticles

The *in vitro* drug release was studied by UV-visible spectrophotometry ^9^. In brief, 40 mg of AT1-HSA-MRN-NPs were suspended in 10 mL of PBS at 37°C and 120 rpm in a shaking incubator. At predetermined time intervals of 0, 0.25, 0.5, 0.75, 1, 2, 4, 8, 18 and 24 hours, 0.5 mL of the nanoparticle suspension was withdrawn and re-substituted with 0.5 mL of fresh PBS. The withdrawn suspension was centrifuged using Amicon centrifugal filters (10K MWCO) and supernatant was used to determine the amount of MRN released. The MRN was detected at 356 nm using a colorimetric assay and a cumulative MRN release over time was calculated ^21^.

### Cell culture and viability

Rat cardiomyoblasts (H9c2) cells were received as a kind gift from Dr. Renzo Cecere, M.D. (Montreal General Hospital, QC, Canada). The H9c2 cells were grown in Dulbecco’s Modified Eagle Medium (DMEM) supplemented with 10% fetal bovine serum (FBS). H9c2 cells were seeded at an initial density of 5000 cells/well in clear bottom 96-well black plates for 24 hours in a humidified incubator at 37°C and 5% CO_2_, Cells were treated with AT1-HSA-MRN-NPs, AT1-HSA-NPs, MRN-HSA-NPs and MRN Lactate, diluted in serum-free cell culture medium. The MRN concentration in the nanoparticles and MRN Lactate were 1 mM, as optimized from previous studies ^12^. After 4, 24 and 48 hours of incubation, the cells were washed thrice with PBS. Cells were treated with 100 μL of fresh cell culture medium and 20 μL of MTT reagent and incubated at 37°C and 5% CO_2_ for 4 hours. The media was removed, and cells were lysed by addition of 100 μL of DMSO for 15 minutes at room temperature. The absorbance was measured at 570 nm using the Victor3V 1420 Multilabel Counter spectrophotometer.

### Overexpression of the AT1 Receptor

The H9c2 cells were seeded at an initial density of 5,000 cell/well in 96-well plates, separated into three groups: Normal, Hypoxic and Hypertrophic. Hypoxia was induced by treatment with 100 μM of CoCl_2_.6H_2_O for 24 hours, which simulated the conditions of MI ^34^. Hypertrophy was induced by treatment with 20 μM H_2_O_2_ for 48 hours to simulate conditions of CHF ^35^. The cells in each group were treated with the anti-AT1 antibody (Abcam, Canada) for 1 hour followed by a goat polyclonal secondary antibody conjugated to Alexa 488 for an additional 1 hour. The cells were fixed with 4% paraformaldehyde in PBS for 10 mins and thrice washed with PBS and stored at 4°C. The fluorescence was measured at 495 nm excitation/ 519 nm emission wavelengths.

### Intracellular Nanoparticle Uptake

The H9c2 cells were seeded at an initial density of 5,000 cell/well in 96-well plates, separated into three groups: Normal, Hypoxic and Hypertrophic. Cells were subjected to hypoxia by treatment with 100 μM of CoCl_2_.6H_2_O for 24 hours, to simulated MI ^34^. Cells were subjected to hypertrophy by treatment with 20 μM H_2_O_2_ for 48 hours to simulate HF ^35^. The H9c2 cells in each group were treated with 0.5 mg/mL of fluorescently tagged AT1-HSA-MRN-NPs, Scr-HSA-MRN-NPs and MRN-HSA-NPs for 4 hours. The Scr peptide was the same amino acid chain as AT1 peptide but in a scrambled order ^20^. The Scr peptide-tagged NPs were used to confirm the targeting efficiency of the AT1 peptide-tagged NPs. The cells were washed thrice with PBS and fresh media was added. The fluorescence intensity was measured at 489nm/535nm using a Victor3V 1420 Multilabel Counter spectrophotometer (Perkin Elmer, Woodbridge, ON, Canada).

## Results

### Nuclear Magnetic Resonance

The design and synthesis of the AT1-HSA was initially characterized by ^1^H-NMR (**Fig. 2**). The NMR spectrum of AT1 displays peaks at around 6.8 ppm which are due to the presence of tyrosine residues, while most of the downfield resonance is due to histidine. This signal is also visible on the AT1-HSA spectrum. The spectrum contains two sets of signals. The more intense signals arise from the trans peptide bonds whereas the less intense signals are due to a cis-bond between proline and histidine ^21^. Peaks at around 4.0 ppm due to glycine residues visible on AT1 spectrum is also visible on the AT1-HSA spectrum. However, the NMR spectra of the HSA-AT1 and HSA in comparison to AT1 peptide displayed broader peaks, which results from the higher molecular weight (66500 Da) of the HSA in comparison with the AT1 peptide (1274 Da). Thus, the data gathered from ^1^H-NMR indicate the conjugation of the AT1 peptide with the HSA, which was also confirmed with Mass Spectrometry.

**Figure 2.**
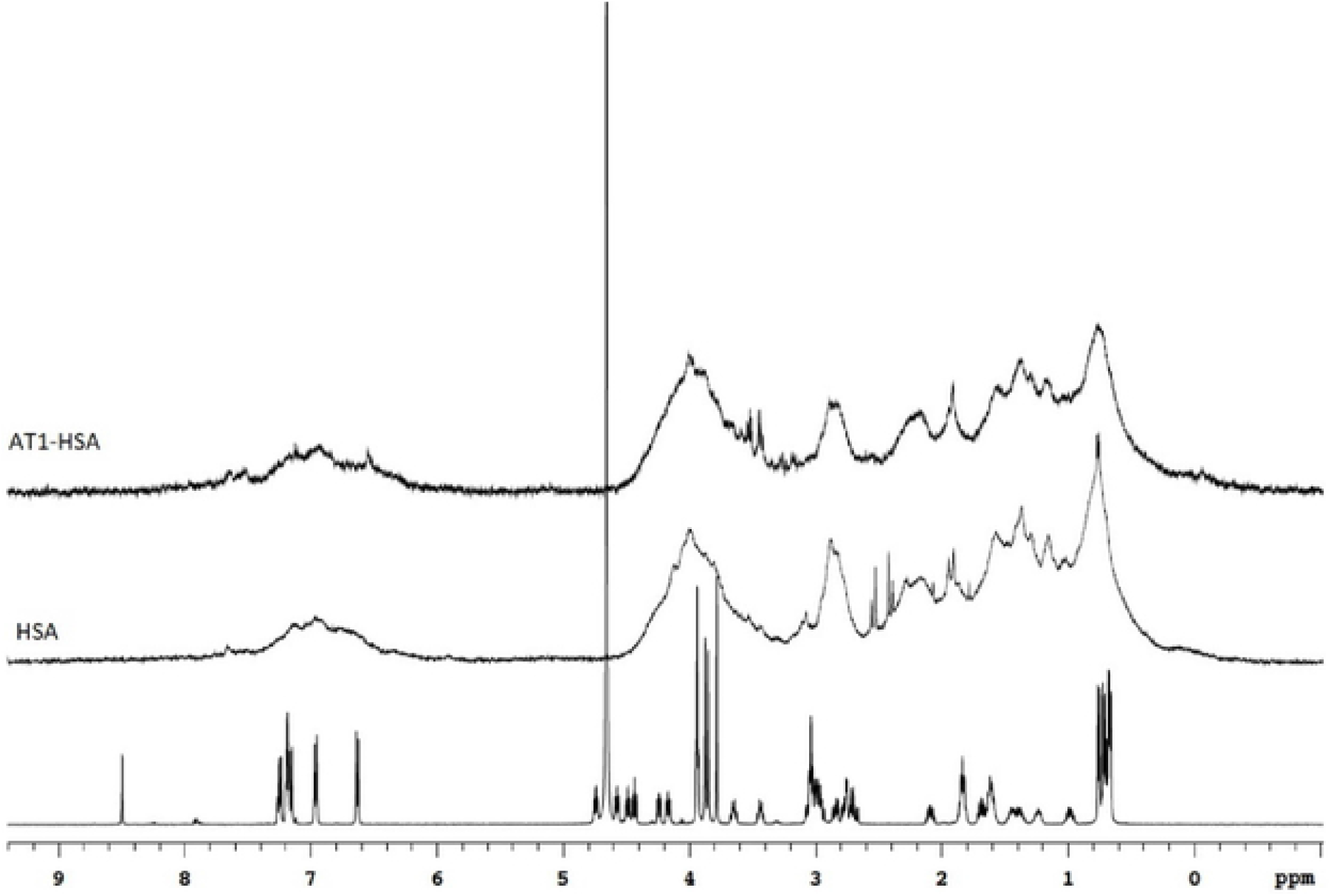
IH NMR based characterization of AT1-HSA, HSA and AT1 peptide. The AT1 peptide exhibits a spectrum with sharp peaks at δ= 0.5-1.0 , δ=1. 1-2. 1, δ= 2.8-3.8 , δ=4 .0-5 .0 and δ=6.3-7.3. The peaks at δ=6.8 due to tyrosine and around δ=3.8-4.0 due to glycine from the ATI peptide can be seen on the ATI-HAS spectrum around δ=3.5-4.0 and 6.8 ppm.

### Mass Spectrometry

To validate the conjugation of AT1 peptide with HSA, MALDI-TOF-MS was used to compare the average molecular weight change between HSA and AT1-HSA. The mass-to-charge ratio (m/z) of the green peak (AT1-HSA) was approximately 7000 higher than the red peak (HSA). The molecular weights of the AT1 peptide, 5(6)-Carboxyfluorescein-NHS, EDC and Sulfo-NHS is 1274, 376.32, 190 and 217 g/mol, respectively. Upon calculating the cumulative molecular weight of AT1-HSA, and comparing with HSA protein, it can be inferred that at least 3 molecules of AT1 were successfully conjugated with the surface of each HSA molecule (**Fig. 3**).

**Figure 3.**
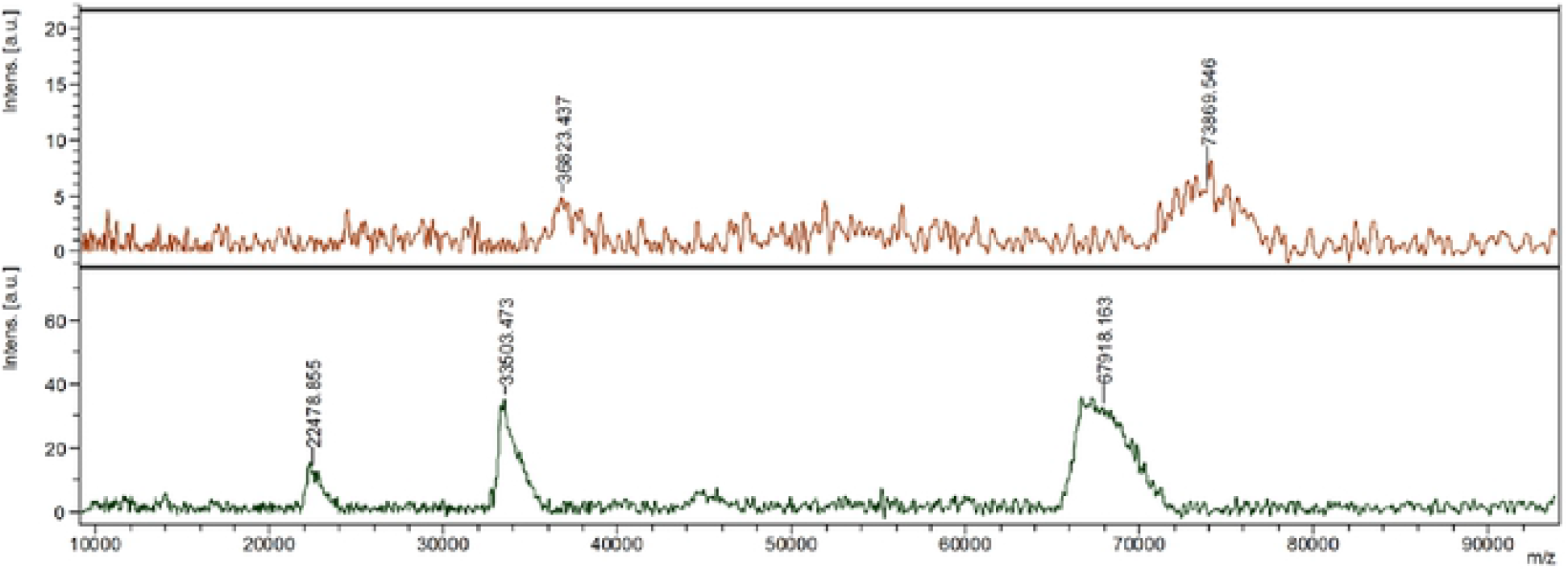
HSA (lower) and AT1-HSA (upper),was analyzed by Matrix-Assisted Laser Desorption/Ionization Time of Flight Mass Spectrometry. The m/z ratio of the AT1-HSA peak was at least 7000 higher than that of the HSA peak, which demonstrated that AT1 was successfully conjugated to the surface of HSA.

### Nanoparticle characterization

The nanoparticles size was determined using DLS and laser Doppler anemometry for zeta potential analysis. The particle size of AT1-HSA-MRN-NPs was 215.2±4.7 nm with a zeta potential of -28.8±2.7 mV, and size of MRN-HSA-NPs (without AT1) was 189.6±3.8 nm with zeta potential of -27.5±4.6 mV. The morphology of the nanoparticles as observed using TEM under 13500X **(Fig. 4(a))** and 55,000X magnification exhibited a near spherical shape with moderately uniform particle size and even distribution **(Fig. 4(b))**. Under 250,000X magnification, the AT1-HSA-MRN-NPs had a dark core surrounded by a bright membrane, which confirmed the distinct layer **(Fig. 4(c))**. The yield of the AT1-MRN-HSA-NPs was 75.6±2.5%, and encapsulation efficiency was 40.5±1.5%.

**Figure 4.**
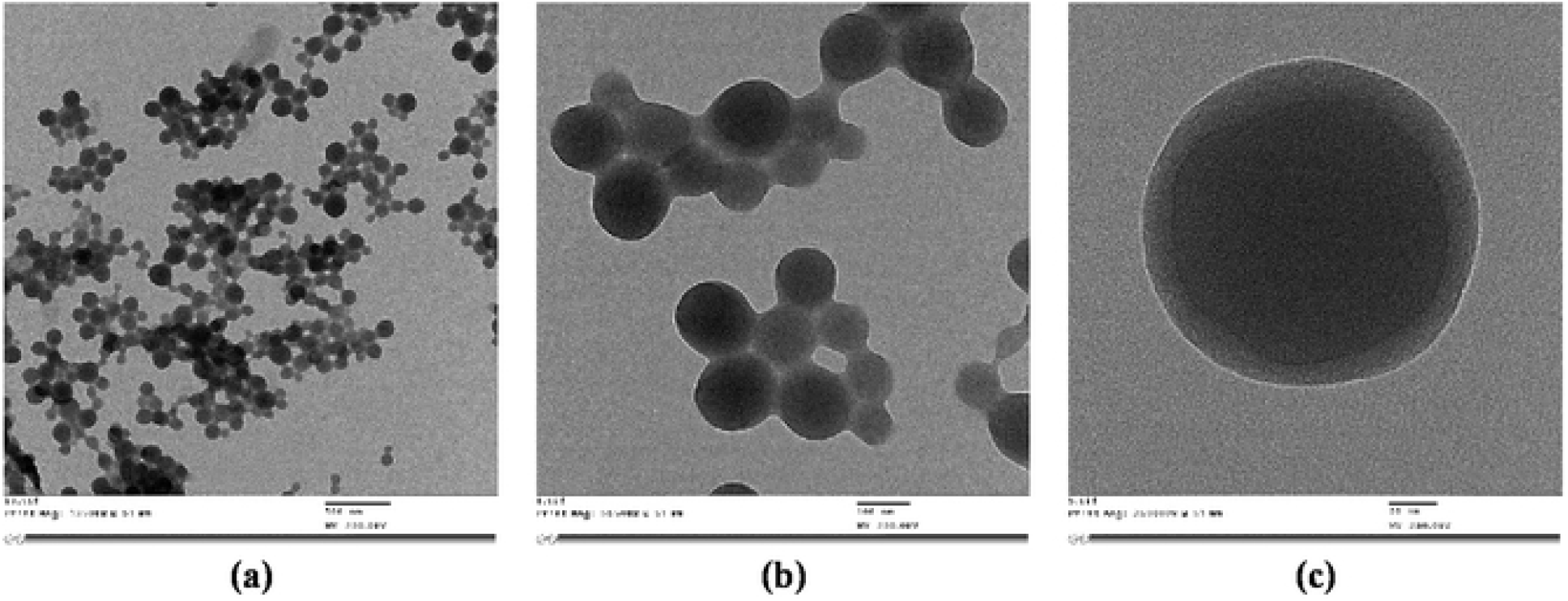
Nanoparticle surface characterization using TEM analysis (a) Under magnification of 13,500X, ATI-HSA-MRN-NPs of size 215.2±4.7 nm and zeta potential of -28.8±2.7 mV (Scale = 500 nm) (b) Under magnification of 55,000X, ATI-HSA-MRN-NPs with moderately uniform particle size (Scale = 100 nm); (c) Under 250,000X magnification, the ATI-HSA-MRN-NPs display a dark core surrounded by a bright membrane, which confirmed the distinct peptide layer (Scale = 20 nm).

### In Vitro Drug Release Study

The drug release from AT1-HSA-MRN-NPs was studied by suspending 40 mg of nanoparticles in 10 mL PBS at 37 °C and 120 rpm, with a starting MRN concentration of 0.8 mg/mL prepared from MRN/AT1-HSA (wt./wt.) ratio of 1:10. MRN showed a sustained release with approximately 50% of the MRN released between 4-6 hrs after which the release became slower with up to 75% of the MRN being released by 24 hrs **(Fig. 5)**.

**Figure 5.**
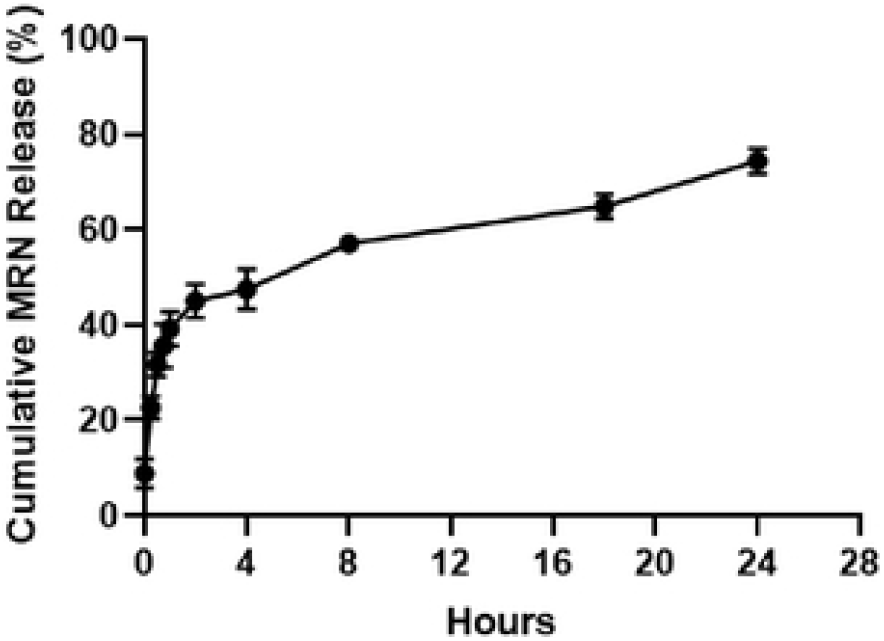
Cumulative drug release (mean ± SD %, n = 3) ofMRN from AT1-HSA-MRN-NPs with starting MRN concentration of 0.8 mg/mL over predetermined time intervals of 0, 0.25, 0.5, 0.75, I, 2 , 4, 8, 18 and 24 hrs.

### Cell Viability Analysis

The MTT assay was performed for cell viability where H9c2 cells were treated with AT1-HSA-MRN-NPs, AT1-HSA-NPs, MRN-HSA-NPs and MRN Lactate containing 1mM MRN concentrations for 4, 24 and 48 hours. The 1mM MRN concentration was optimized in our previous study and is being used here for consistency of results ^33^. The safety analysis of MRN-HSA-NPs was conducted using HUVEC and H9c2 cells in our previous study and therefore was not repeated here ^33^. Results showed that the H9c2 cells treated with AT1-HSA-MRN-NPs, AT1-HSA-NPs and MRN Lactate for 4 hours showed cell viability of 73.4±1.4%, 101.7±2.9% and 44.9±1.4%, respectively **(Fig. 6(a))**. At 24 hours, the cell viability was 55.5±3.7%, 70.8±11.2% and 41.2±3.8%, respectively **(Fig. 6(b))**, and at 48 hours, 52.4±2.1%, 65.9±11.2% and 35.3±7.7%, respectively **(Fig. 6(c))**. Therefore, the AT1-HSA-MRN-NPs exhibit better safety characteristics and lesser cytotoxicity as compared to MRN Lactate. The *in vitro* cytotoxicity due to hypoxia and hypertrophy treatments was also investigated and compared with normal H9c2 cells. No significant difference in cell viability was observed between the 3 conditions **(Fig. 7)**.

**Figure 6.**
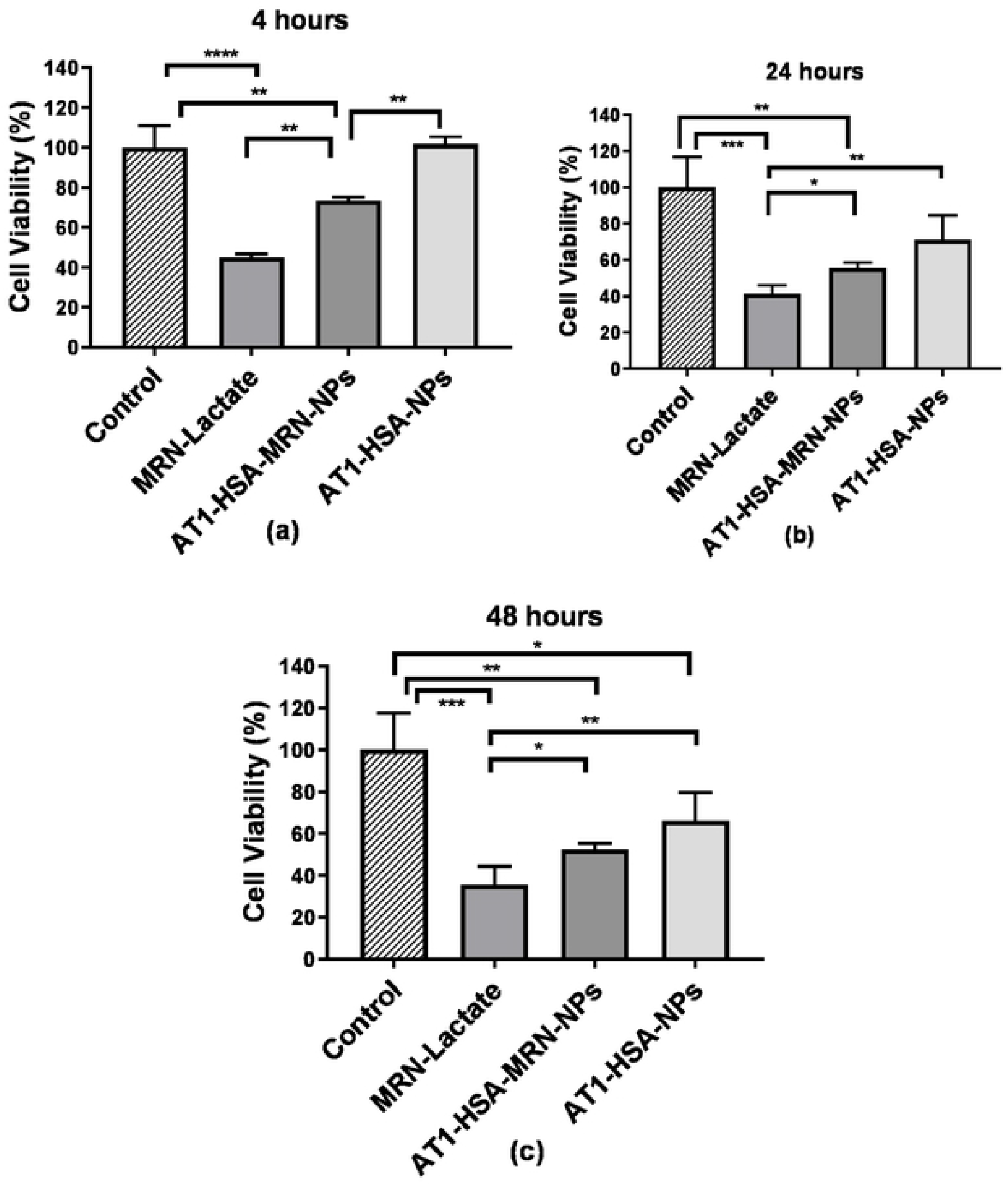
Cell viability analysis on H9c2 cells treated ,vith MRN-Lactatc, AT1-HSA-MRN-NPs and AT1-HSA-NPs at 1 mM MRN concentration at (a) 4 hours, (b) 24 hours and (c) 48 hours. The graph shows a representative result of mean ± SD (n=5). \*\*\*\**P*<0.0001 was considered highly significant and \*\*\**P*< 0.001, \*\**P*<0.01 and \**P*<0 .05 were considered significant based on Tukey’s posthoc analysis.

**Figure 7.**
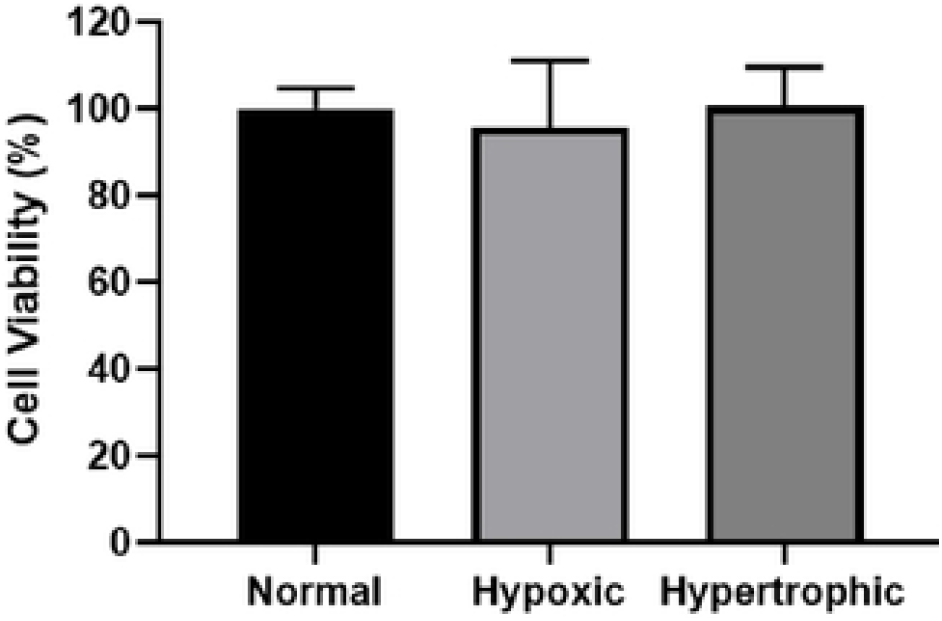
Viability of H9c2 cells treated for hypoxia and hypertrophy in comparison with normal cells (non-hypoxic and non-hypertrophic). There were no significant differences *(P*>0. l) in cell viability amongst the groups.

### Angiotensin II Type 1 Receptor Overexpression

The AT1R receptor overexpression was studied in hypoxic, hypertrophic and normal H9c2 cells. Results showed that hypoxic and hypertrophic cells exhibited a significantly higher expression of the AT1 receptors as compared to normal cells (P<0.0001). The fluorescence intensity displayed by the hypoxic and hypertrophic cells was almost twice higher than that observed in the normal cells **(Fig. 8)**.

**Figure 8.**
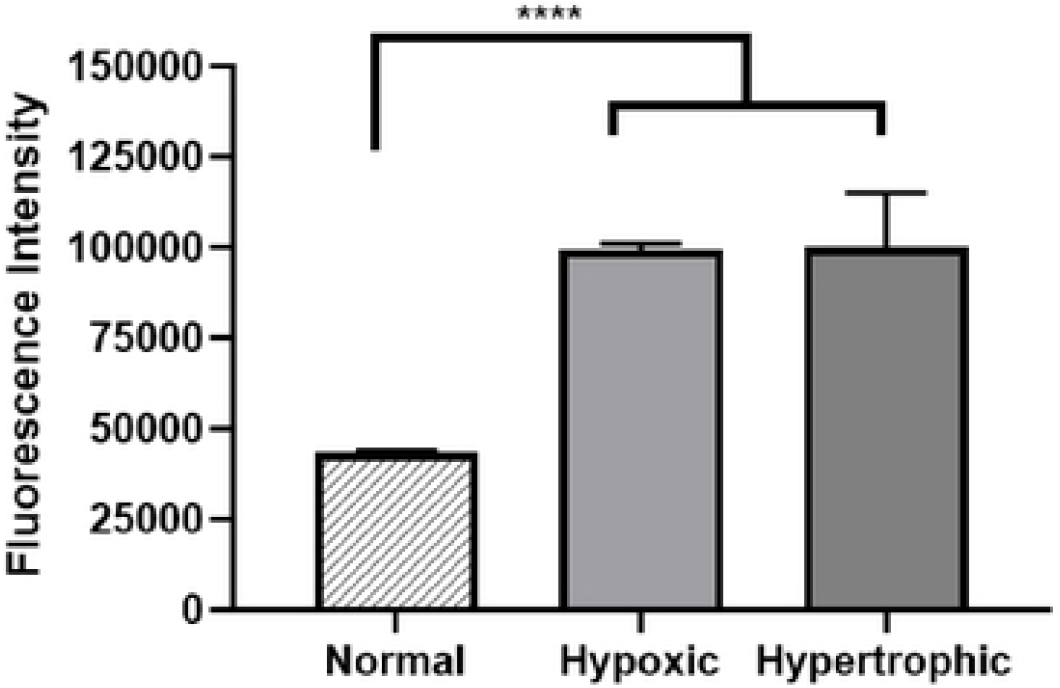
AT1 expression in normal, hypoxic and hypertrophic H9c2 cells. Results are represented as mean ± SD (n=5). \*\*\*\**P*<0.0001 was considered highly significant based on Tukey’s posthoc analysis.

### Intracellular Nanoparticle Uptake Analysis

Since AT1 receptors are overexpressed on cardiomyocytes during HF, conjugating the nanoparticles with the AT1 peptide was anticipated to demonstrate higher uptake of the AT1-HSA-MRN-NPs through receptor-mediated endocytosis. The nanoparticle concentration was 0.5 mg/mL. In normal (non-hypoxic, non-hypertrophic) cells, the AT1-HSA-MRN-NPs exhibited significantly higher fluorescence intensity as compared to the non-targeted MRN-HSA-NPs (P<0.001) and Scr-HSA-MRN-NPs (P<0.01) **(Fig. 9(a))**. Similarly, in hypoxic cells, the AT1-HSA-MRN-NPs uptake was significantly higher (P<0.0001) than that of MRN-HSA-NPs and Scr-HSA-MRN-NPs **(Fig. 9(b))**. Also, for hypertrophic cells, the uptake of AT1-HSA-MRN-NPs was significantly greater (P<0.0001) than that of MRN-HSA-NPs and Scr-HSA-MRN-NPs **(Fig. 9(c))**.

**Figure 9.**
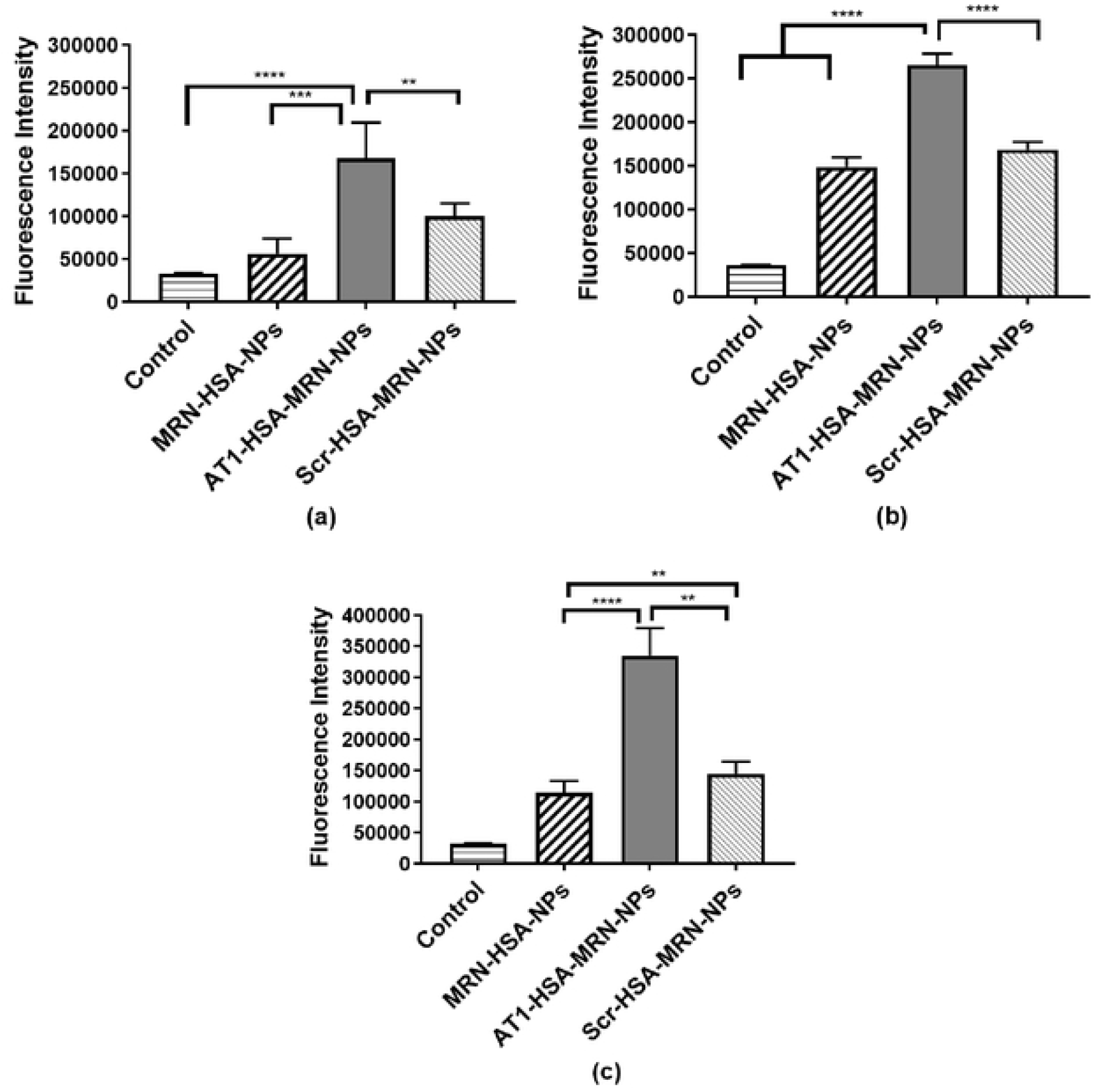
Intracellular uptake of MRN-HSA-NPs, AT1-HSA-MRN-NPs and Scr-HAS-MRN-NPs in H9c2 cells (a) Normal cells (non-hypoxic, non-hype rtrophic), (b) Hypoxic cells and (c) Hypertrophic cells. The nanoparticle concentration was 0.5 mg/mL. The graph shows a representative result of mean ± SD (n=5). \*\*\*\**P*<0.0001 was considered highly significant and \*\*\**P*<0.001, \*\**P*<0.01 were considered significant based on Tukey’s posthoc analysis.

Also, H9c2 cells under hypoxia and hypertrophy exhibited greater uptake of AT1-HSA-MRN-NPs as compared to normal conditions however, the uptake of MRN-HSA-NPs and Scr-HSA-MRN-NPs was significantly less (\*\*\**P<*0.001).

## Discussion

The urgent need to effectively treat MI and HF has led to research and innovation of various new strategies and treatment modalities. These include delivery of drugs, growth factors, cytokines and other molecules for myocardial regeneration or treatment ^36,37^.

However, due to inherent limitations with most treatment strategies such as lack of target specificity, low bioavailability, cardiac rejection while heart pumping, or non-specific distribution, the therapeutic effect is lessened ^38^. The emerging studies on nanoparticles and targeted drug delivery systems have displayed promising results, however, their efficacy remains dependent on the drug binding capacity, solubility, nanoparticle degradability and plasma retention time ^39^. In this study, keeping in view the intended features of ideal drug delivery systems, a targeted nanoparticle formulation was synthesized. The HSA surface was modified to attach a targeting ligand, AT1 peptide, to achieve superior delivery characteristics. The AT1 peptide shows specificity for the AT1 receptor present on the myocardium, which is found to be overexpressed under CHF conditions ^6-8^. Using the AT1 peptide as the targeting moiety will facilitate receptor-mediated nanoparticle uptake.

The AT1 peptide was conjugated to the HSA surface through a two-step covalent chemical reaction. The 5(6)-Carboxyfluorescein-NHS targets primary amines such as in the side chain of lysine residues, to form stable amide bonds. This allows the carboxylic group to undergo a carbodiimide reaction with EDC, at pH 5.5, forming an unstable amine-reactive O-acylisourea intermediate. This unstable intermediate is further reacted with the Sulfo-NHS and the amine groups on AT1 peptide to release urea as a byproduct and form stable AT1-HSA. The pre-modification of the HSA by conjugation with the AT1 peptide was preferred over post-modification of the HSA-NPs as the latter may cause drug loss and leakage during the synthesis and purification. NMR and MALDI-TOF confirmed the binding of AT1 peptide to the HSA surface. In NMR results, HSA shows broad peaks owing to the high molecular weight of 66.5 kDa, whereas the AT1 peptide, with a molecular weight of ∼ 1274 g/mol exhibits a clear spectrum showing chemical shifts due to the presence of the amino acid residues. Due to the large differences in molecular weights, the AT1-HSA spectrum resembles the HSA with visible differences at 6.8 and 4.0 ppm indicating AT1 binding. To confirm this, MALDI-TOF-MS was performed. The spectrum on the x-axis represents the m/z (mass/charge) ratio which was 73869.546 for AT1-HSA and 67918.163 for HSA, which was 7000 higher. Considering the mass of the AT1 peptide, the HSA and the cross-linkers, at least 3 AT1 peptide molecules were bound to the HSA surface.

The AT1-HSA-MRN-NPs were formed by the ethanol desolvation process ^12,30^. The nanoparticles exhibited a spherical structure as observed under TEM. The AT1-HSA-MRN-NP size was 215.2±4.7 nm with a zeta potential of -28.8±2.7 mV, in comparison with AT1-HSA-NPs with a size of 189.6±3.8 nm and zeta potential of -27.5±4.6 mV. The nanoparticles were less than 250 nm with a negative zeta potential, which suggests greater physical stability as nanoparticle aggregation is prevented due to presence of negative charges ^40^. The particle size being less than 250 nm indicates a prolonged blood circulation time as the particles are not removed easily through opsonization ^41,42^. MRN release over 24 hrs from the nanoparticles further confirmed drug binding to the AT1-HSA-MRN-NPs. The binding characterization of MRN with HSA has been studied extensively in our previous study and was therefore not repeated here ^33^.

The *in vitro* cytotoxicity was evaluated on H9c2 cells treated with MRN-Lactate, AT1-HSA-MRN-NPs and AT1-HSA-NPs. MRN is clinically administered as a lactate formulation intravenously to adult as well as pediatric patients for HF and associated cardiac conditions ^43^. However, the use of MRN Lactate is linked with side effects such as palpitation, cardiac arrythmia and renal dysfunction ^44^. This may be attributed to the non-targeted delivery of MRN Lactate and hence the need for a continuous infusion to meet the dosage requirements. Using the targeted nanoparticle formulation, synthesized in this study, as drug carriers this limitation would be overcome, given their higher biocompatibility, higher retention time, drug binding capacity and characteristics of controlled drug release ^45^. The cell viability of H9c2 cells was investigated at 4, 24 and 48 hours, with MRN Lactate exhibiting higher cytotoxicity as compared to AT1-HSA-MRN-NPs and AT1-HSA-NPs. Nanoparticle safety is essential as to ensure their suitability for use in future pre-clinical and clinical studies.

The intracellular uptake of the nanoparticles was investigated in normal (non-hypoxic, non-hypertrophic), hypoxic and hypertrophic H9c2 cells and H9c2 cells have been found to be more suitable for cardiac ischemia studies ^46^. Inducing hypoxia and hypertrophy in cells closely mimics MI and HF conditions ^35,47,48^. The cell viability analysis comparing the normal cell viability with that of hypoxic and hypertrophic H9c2 cells suggested that the hypoxia and hypertrophy inducing treatments were safe and did not cause cytotoxicity. Literature has widely suggested that under HF, the AT1 receptors present on the cardiomyocytes are overexpressed and these receptors could be blocked to reverse cardiac remodeling ^7,8^. Therefore, targeting the overexpressed AT1 receptors with the targeted AT1-HSA-MRN-NPs to hypoxic and hypertrophic cardiac cells allowed higher uptake of the AT1-HSA-MRN-NPs as compared to the non-targeted MRN-HSA-NPs and Scr-HSA-MRN-NPs. The Scr peptide-tagged nanoparticles were used as a negative control to ensure that the higher uptake of AT1-HSA-MRN-NPs was a direct result of AT1 peptide mediated targeting and not passive uptake ^5^. Also, the uptake of the AT1-HSA-MRN-NPs was significantly higher in hypoxic and hypertrophic cells vs the normal cells. These studies confirm the targeting ability of the AT1 peptide under MI and HF conditions and demonstrate that AT1-HSA-MRN-NPs can safely be used *in vivo* as targeted drug delivery systems for congestive heart failure and related conditions.

## Conclusion

To summarize, we have synthesized and developed stable AT1 peptide conjugated albumin nanoparticles to deliver MRN in a targeted manner for heart failure treatment. This novel drug delivery system demonstrates physical stability, release, biocompatibility, specific targeting ability and higher cellular uptake. Also, as compared to the non-targeted MRN Lactate, AT1-HSA-MRN-NPs show greater biocompatibility in cardiomyoblasts. In future, the performance of AT1-HSA-MRN-NPs will be evaluated in a rat model of CHF along with MRN pharmacokinetics. This targeted therapy is anticipated to be more effective in improve the cardiac function in CHF as compared to the currently available treatments.

## Acknowledgements

This work is supported by the research funding granted to Dr. Satya Prakash from Canadian Institute of Health Research (CIHR) and the Natural Sciences and Engineering Research Council (NSERC). The authors are grateful to Mr. Xue Dong Liu for assistance in TEM imaging (Facility for Electron Microscopy Research, Materials Engineering, McGill University), Dr Tara Sprules for assistance with NMR and Mr. Nadim Saadeh for help with Mass Spectroscopy (Mass Spectroscopy Facility, Department of Chemistry, McGill University).

## Disclosure

The author reports no conflicts of interest in this work.

## Data Availability Statement

The authors confirm that the data supporting the findings of this study are available within the article [and/or] its supplementary materials.

## Notes

### Competing Interest Statement

The authors have declared no competing interest.

